# Fluorescence lifetime imaging of pH along the secretory pathway

**DOI:** 10.1101/2021.03.29.437519

**Authors:** Peter T.A. Linders, Melina Ioannidis, Martin ter Beest, Geert van den Bogaart

## Abstract

Many cellular processes are dependent on correct pH levels, and this is especially important for the secretory pathway. Defects in pH homeostasis in distinct organelles cause a wide range of diseases, including disorders of glycosylation and lysosomal storage diseases. Ratiometric imaging of the pH-sensitive mutant of green fluorescent protein (GFP), pHLuorin, has allowed for targeted pH measurements in various organelles, but the required sequential image acquisition is intrinsically slow and therefore the temporal resolution is unsuitable to follow the rapid transit of cargo between organelles. We therefore applied fluorescence lifetime imaging microscopy (FLIM) to measure intraorganellar pH with just a single excitation wavelength. We first validated this method by confirming the pH in multiple compartments along the secretory pathway and compared the pH values obtained by the FLIM-based measurements with those obtained by conventional ratiometric imaging. Then, we analyzed the dynamic pH changes within cells treated with Bafilomycin A1, to block the vesicular ATPase, and Brefeldin A, to block ER-Golgi trafficking. Finally, we followed the pH changes of newly-synthesized molecules of the inflammatory cytokine tumor necrosis factor (TNF)-α while they were in transit from the endoplasmic reticulum via the Golgi to the plasma membrane. The toolbox we present here can be applied to measure intracellular pH with high spatial and temporal resolution, and can be used to assess organellar pH in disease models.

## Introduction

Physiological pH homeostasis is crucial for many cellular processes. Not only the cytosolic pH is of importance, but defined intraorganellar pH delineates the secretory pathway. The pH of the endoplasmic reticulum (ER) is approximately 7, while the Golgi apparatus slightly acidifies from pH 6.7 at the *cis* face to pH 6.0 at the *trans* face^1–3^. Before secretory cargo is released at the plasma membrane and reaches the neutral pH of the extracellular environment, the pH in secretory vesicles is about 5.2^1,2^.

pH is not only crucial for proper protein folding and enzyme activity through influencing the charge of amino acid side chains, but its importance in secretory protein transport is increasingly clear^4^. pH affects binding affinities of cargo molecules to trafficking chaperones, and thereby pH differences facilitate intracellular transport by both influencing the transit of cargo^5–11^ and the sorting of secretory pathway resident proteins^12–14^. Moreover, the localization of glycosylation enzymes and their substrates is determined by pH^4,15–18^, and defects in this homeostasis cause a wide range of human disease^4,19–25^. Being able to accurately determine intraorganellar pH along the secretory pathway is therefore of both fundamental and diagnostic importance.

Fluorescent dyes that allow the measurement of intraorganellar pH exist and are commercially available^26–30^, but the inability of specific organellar targeting is a major drawback. The pH in the lumen of the Golgi and ER in mammalian cells have been measured using Shiga-like toxins covalently bound to fluorescent dyes^31,32^ and with the biotin-avidin system^33^. However, especially the development of pH-sensitive mutants of green fluorescent protein (GFP), such as pHLuorin^34,35^, which can be targeted to specific organelles by fusion proteins, have enabled specific measurement of intracellular compartments. Two classes of pHLuorin were developed by mutagenesis which altered the bimodal excitation spectrum of GFP with peaks at 395 and 475 nm^34,36^. First, ecliptic pHLuorin, which shows a reduction of its excitation efficiency at 475 nm at pH values lower than 6. Second, ratiometric pHLuorin which shows a gradual increase in the ratio of excitation at 475/395 nm between pH 5.5 and pH 7.5^34^. With ecliptic pHLuorin, intraorganellar pH can be determined by first recording an image at 475 nm excitation, and then correlating the fluorescence intensities with a calibration curve. The pH can be determined with ratiometric pHLuorin using a similar approach, but now by sequentially recording images at 395 and 475 nm excitation. A new version of ratiometric pHLuorin, ratiometric pHLuorin2 (RpHLuorin2) was later developed with 8-fold improved fluorescence^35^.

Ecliptic pHLuorin is less accurate than ratiometric pHLuorin, because the fluorescence intensity not only depends on the pH but also on the concentration of pHLuorin. However, ratiometric imaging also has several drawbacks, such as sensitivity to background fluorescence leading to high variation in the ratio values and the need for two sequential image acquisitions with two different excitation wavelengths. As the exocytic pathway is highly dynamic, the sequential imaging could potentially result in misalignment of the emitted signal, compromising the calculation of ratio values. To overcome this problem of dual excitation, GFP-based probes have been developed that show a pH-dependent change in fluorescence emission, including E^2^GFP^37^ and deGFP4^38,39^, and pH-sensitive fluorophores have been targeted to organelles with the HaloTag technology^40^. However, spectral overlap in fluorescence emission wavelengths limits the use of these probes for multi-color experiments together with other fluorescent probes.

In this study, we took another approach to overcome the problem of dual excitation and exploit fluorescence lifetime, an intrinsic property of fluorophores that is insensitive to changes in laser intensity or protein concentration^41,42^ but is sensitive to pH^43,44^, to accurately measure intraorganellar pH with both high spatial and temporal resolution.

## Results

### FLIM measurements of recombinant ratiometric pHLuorin2

We first measured the fluorescence excitation spectra of recombinant RpHLuorin2 (Supplementary Figure 1) in different pH solutions with a fluorescence spectrometer. Depending on the pH of the medium, the p-hydroxybenzylidene-imidazolidinone moiety in the chromophore of pHluorin2, a derivative of GFP^35^, can exist in either the neutral phenol form or the anionic phenolate form^45^. As expected^34^, we observed strong dependence of the excitation efficiencies on pH, as a higher pH resulted in an increased emission brightness (at 508 nm) at an excitation wavelength of 470 nm, whereas the fluorescence brightness was reduced at an excitation wavelength of 405 nm (Supplementary Figure 1a). These data show that for pHluorin2, the anionic form shows an excitation peak at 405 nm, while the peak with 470 nm corresponds to the neutral form. We then plotted the ratios of the emission signals with 405 nm over 470 nm excitation as a function of the pH and fitted these data with a dose response relationship, an empirical model to fit the sigmoidal data as the (de)protonation states of RpHLuorin2 will saturate at very high and low pH values (Supplementary Figure 1b). The largest changes in fluorescence of RpHLuorin2 were observed between pH 5.5 and pH 7, making RpHLuorin2 an excellent candidate for pH measurements in the secretory pathway.

As ratiometric determination of pH with RpHLuorin2 requires two sequential image acquisitions with different excitation wavelengths, we investigated whether fluorescence lifetime imaging microscopy (FLIM) would be an appropriate substitute to allow for single-scan imaging. We hypothesized that as the lifetime of fluorophores is influenced by pH^43,44^, the pH sensitivity of RpHLuorin2 would allow for accurate pH measurement based on fluorescence lifetime. Therefore, we performed FLIM of recombinant RpHLuorin2 in different pH solutions at 470 nm excitation (Figure 1). For GFP, the fluorescence lifetime of the phenolate form is in the 2–3 ns range, while that of the phenol form is < 100 ps^46,47^. At 470 nm excitation, we will mainly excite the (fast) phenol form, but (due to fluorescence cross-talk) there will also be some contribution of the (slow) phenolate form. The observed fluorescence lifetime can hence be regarded as a mixture of the decays of the neutral and anionic forms. Thus, an increase in pH will result in a net higher lifetime, as has been reported for other GFP-derived fluorescent proteins^43,48^.

**Figure 1.**
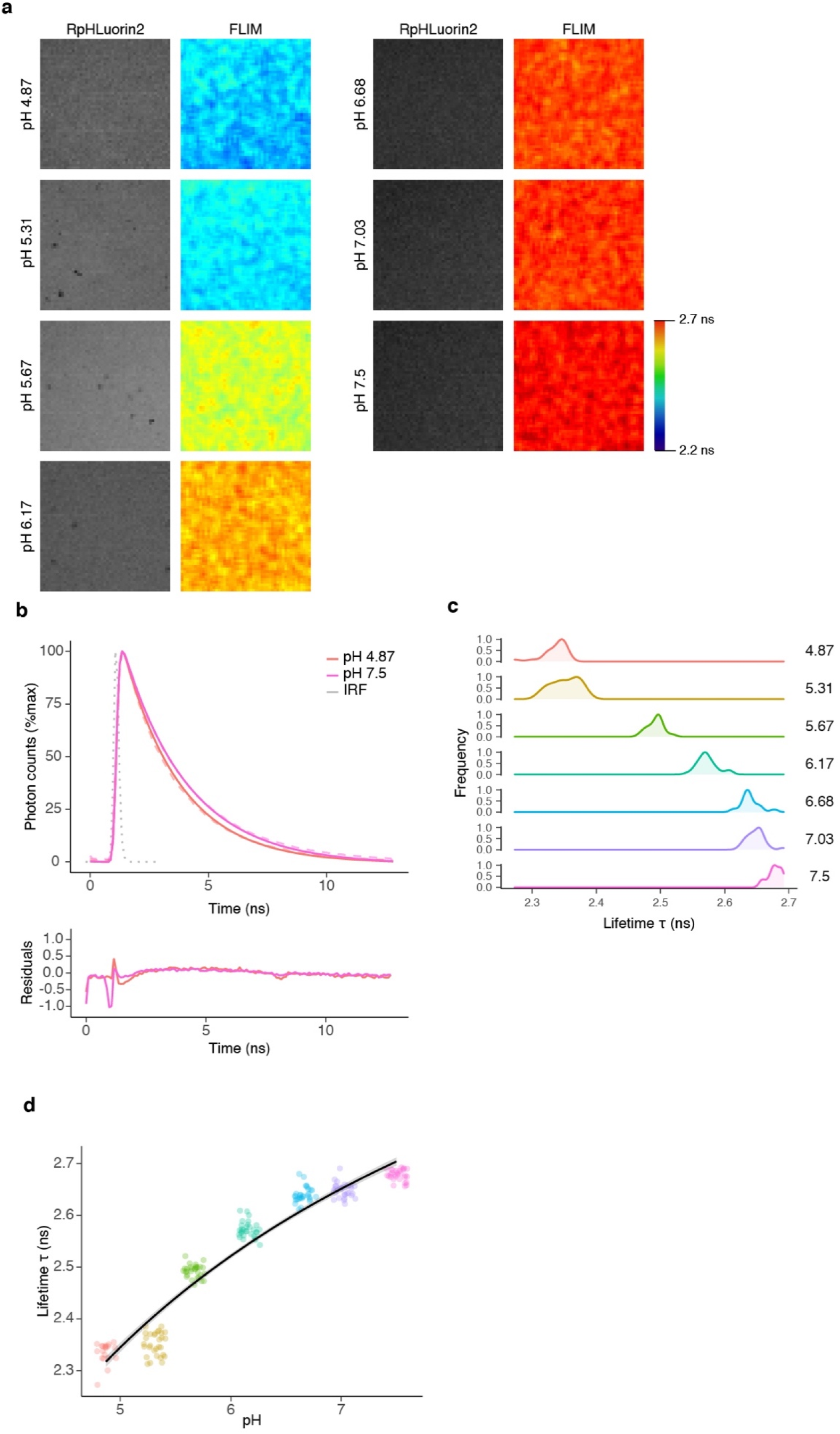
Fluorescence lifetime imaging microscopy (FLIM) of recombinant RpHLuorin2. (a) Representative confocal images of 10 μM recombinant RpHLuorin2 in calibration buffers with defined pH. The intensity image (left column) was convoluted with the fluorescent lifetime value per pixel and pseudo-colored (right column). (b) Representative fluorescence lifetime histograms of recombinant RpHLuorin2 in pH 4.87 solution (red dashed line) or pH 7.5 solution (pink dashed line). Fits with mono-exponential decay functions (pH 4.87, solid red line; pH 7.5, solid pink line) convoluted with the instrumental response function (IRF, gray dotted line). Graphs are normalized to the maximum photon counts. (c) Average lifetime histograms from the images of panel (a). 30 regions of interest were selected per pH buffer and the average lifetime τ was measured. (d) pH dependence of recombinant RpHLuorin2 in defined pH calibration buffers from the images of panel (a).

For purified recombinant pHLuorin2, we observed a dependency of the lifetime as a function of pH, and the fluorescence lifetime increased upon an increasing pH (Figure 1). However, the fluorescence lifetime changed over a larger range of pH values (4.5–7.5; Figure 1d) than the ratio of 405/470 nm excitations (5.5–7.5, Supplementary Figure 1b). This larger dynamic range, which is likely caused by a second protonation event, is an advantage of the FLIM-based approach, because it increases the range of pH values that can be determined. We then fused RpHLuorin2 to several intraorganellar markers in the secretory pathway to perform pH measurements in living cells.

### pH measurements in the secretory pathway

In order to accurately measure intraorganellar pH of specific organelles, we targeted RpHLuorin2 intracellularly by fusing it to proteins and targeting sequences that locate to specific subcellular locations in the secretory pathway (Figure 3a). The pH range that can be measured with lifetime-based measurements of pHLuorin2 (4.5–7.5) is ideally suited for measuring pH along the secretory pathway, as the pH is neutral within the ER (~7), slightly acidic (~6) in the Golgi network, and about 5.2 in secretory vesicles^1–3^. To interrogate the luminal pH along the entire secretory pathway, we fused RpHLuorin2 to the signal sequence of the ER-resident protein calreticulin and a C-terminal ER retention signal KDEL for ER targeting, to the luminal regions of *cis*-/medial-Golgi protein alpha-1,6-mannosyl-glycoprotein 2-beta-N-acetylglucosaminyltransferase (MGAT2), to *trans*-Golgi enzyme beta-1,4-galactosyltransferase 1 (GalT), to *trans*-Golgi network protein TGN46, and to lysosome-associated membrane glycoprotein 1 (LAMP1) for lysosomal targeting, and finally to a GPI anchor for plasma membrane (i.e., extracellular) localization. For the Golgi enzymes (MGAT2 and GalT), we truncated each protein by removing their catalytic sites and only kept the transmembrane region and stalk regions responsible for their localization^49–51^.

We then expressed the fusion constructs in HeLa cells, and recorded FLIM images. We used the GPI-anchored RpHLuorin2 (GPI-RpHLuorin2) to calibrate the probe expressed in cells using the same pH buffers as used for the calibration of purified RpHLuorin2 (Figure 2). We again observed a dependency of the fluorescence lifetime of RpHLuorin2 on pH, although the absolute fluorescence lifetime values were lower than for the recombinant RpHLuorin2, possibly due to crowding effects leading to fluorescence self-quenching^52^. These crowding effects, as well as intracellular pools of GPI-pHLuorin2, likely also contributed to the variation among cells. The fluorescence lifetime dependency on pH could again be fitted with a sigmoidal dose-response model.

**Figure 2.**
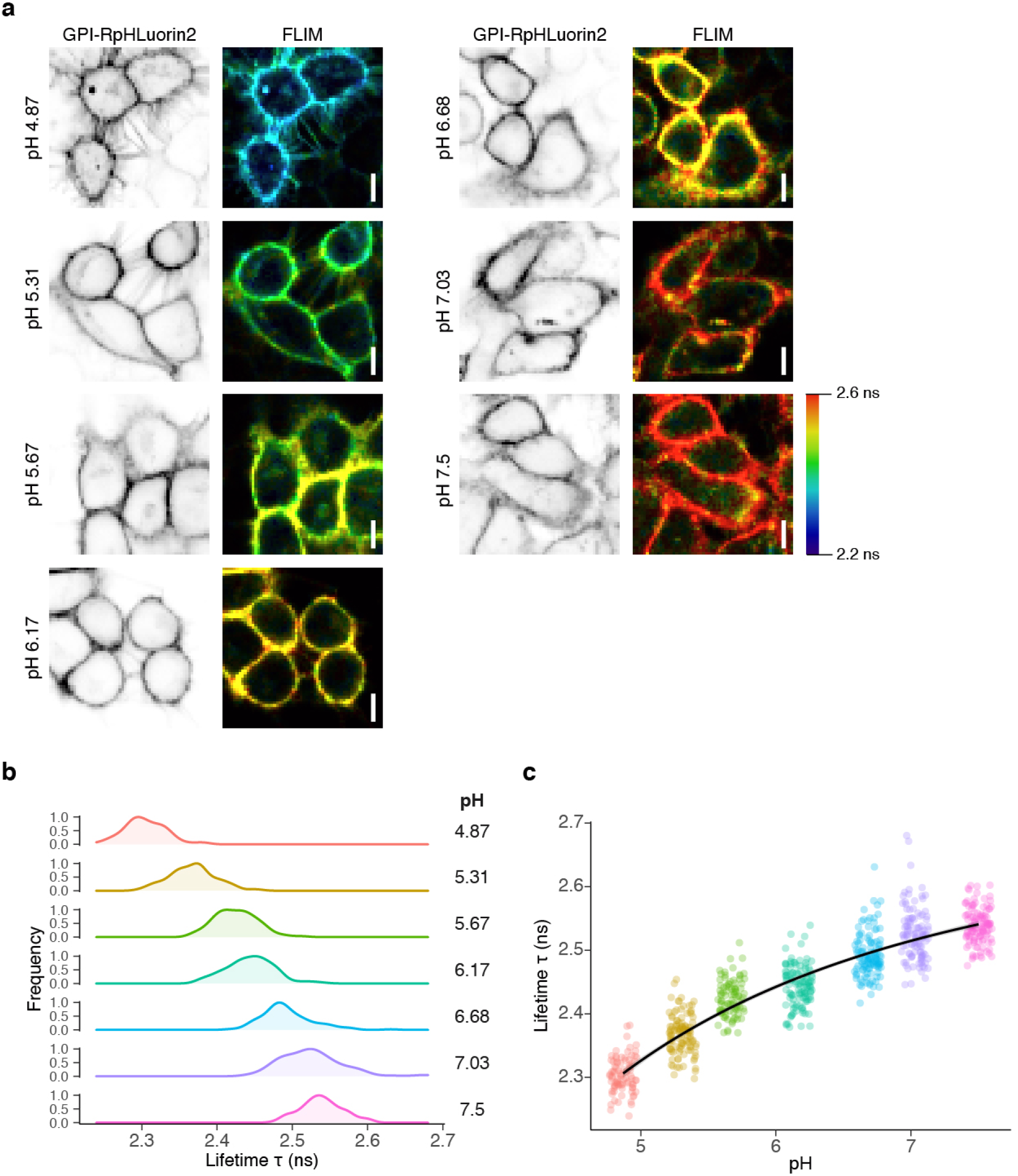
Calibration of RpHLuorin2 by fluorescence lifetime imaging microscopy (FLIM) in HeLa cells expressing GPI-RpHLuorin2. (a) Representative confocal micrographs of HeLa cells expressing GPI-RpHLuorin2 in defined calibration buffers. The intensity image (left column) was convoluted with the fluorescent lifetime value per pixel and pseudo-colored (right column). Scalebars, 10 μm. (b) Average lifetime histograms from the images of panel (a). N = 86 (pH 4.87), 108 (pH 5.31), 90 (pH 5.67), 115 (pH 6.17), 122 (pH 6.68), 113 (pH 7.03) and 120 (pH 7.5) cells from three independent experiments. (c) pH dependence of HeLa cells expressing GPI-RpHLuorin2 in defined pH calibration buffers from the images of panel (a).

After successfully calibrating our system, we proceeded with pH measurements in the lumen of the organelles along of the secretory pathway (Figure 3). With ER-RpHLuorin2, we measured an apparent average pH of 7.2 (95% confidence interval (CI) ± 0.08), while with medial-Golgi marker MGAT2-RpHLuorin2 we measured an apparent average pH of 6.1 (95%CI ± 0.07), and with *trans*-Golgi marker GalT-RpHLuorin2 an apparent average pH of 5.9 (95%CI ± 0.07) (Figure 3b, c). Finally, for lysosomal marker LAMP1-RpHLuorin2 we measured an apparent average pH of 4.7 (95%CI ± 0.15) (Figure 3b, c). These pH values are all consistent with previous literature^1,26^. The fluorescence intensities (numbers of photons collected per cell) did not correlate with the fluorescence lifetimes for the four measured probes (Supplementary Figure 2), indicating that intercellular variations were not caused by differences in expression levels.

**Figure 3.**
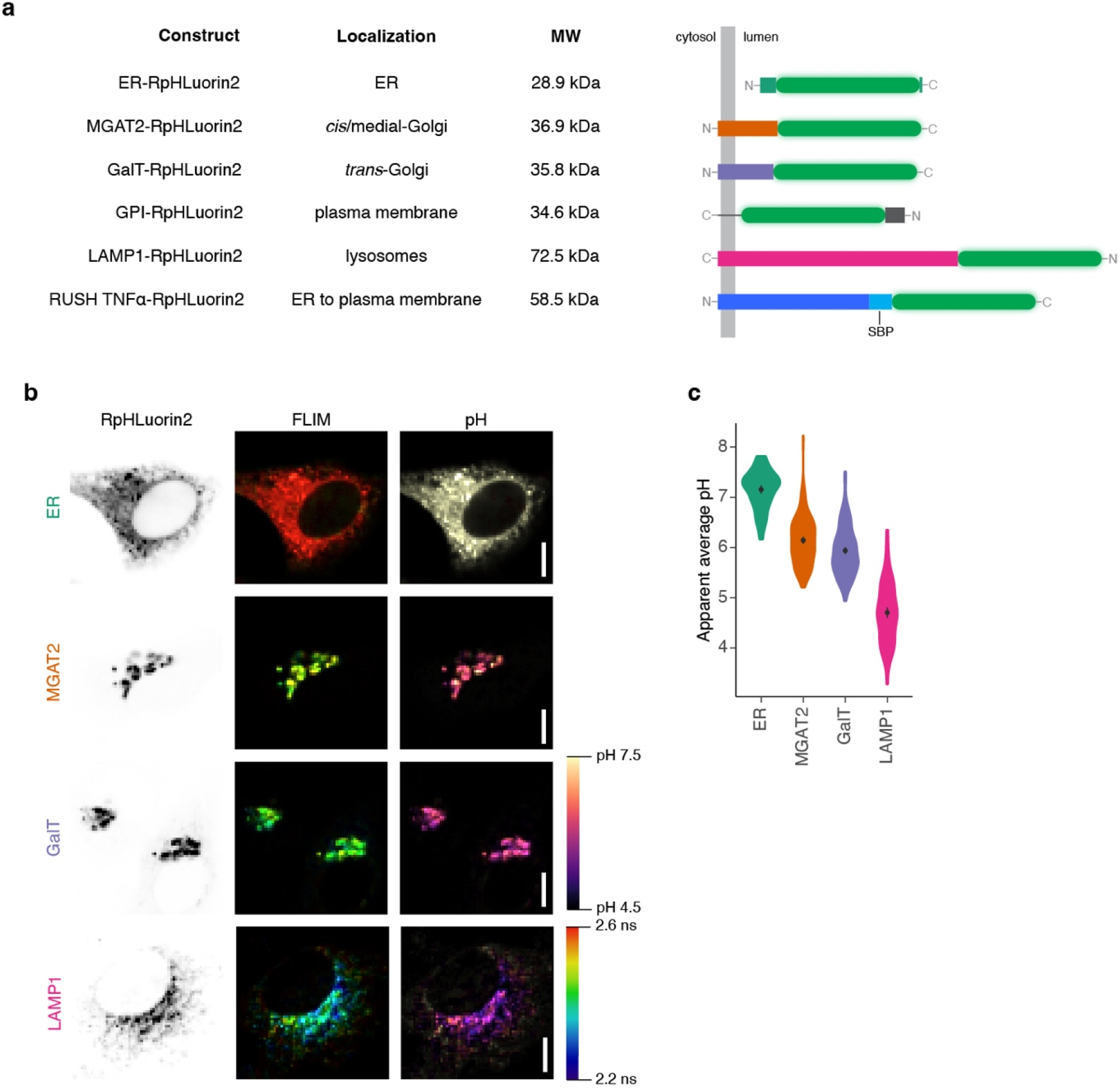
Steady-state pH measurements of secretory pathway markers. (a) Schematic overview of all RpHLuorin2 constructs used in this study. The signal sequence of LAMP1 is removed following co-translational ER insertion and is not shown in the diagram. MW, molecular weight. RUSH, retention using selective hooks^43^. SBP, streptavidin binding protein. (b) Representative confocal micrographs of HeLa cells expressing the mentioned RpHLuorin2 fusion constructs. The intensity image (left column) was convoluted with the fluorescent lifetime value per pixel and pseudo-colored (middle column). The intensity image was also convoluted with the calculated pH per pixel and pseudo-colored (right column). FLIM, fluorescence lifetime imaging microscopy. Scalebars, 10 μm. (c) Quantification of average pH values from panel (b). N = 88 (ER), 188 (MGAT2), 193 (GalT), and 134 (LAMP1) cells from 3 – 5 independent experiments.

To compare the FLIM-based measurements with ratiometric measurements, we also performed ratiometric imaging in cells using confocal laser-scanning microscopy where we changed the excitation wavelength each line of the imaging (Supplementary Figure 3). Confirming our previous experiments with purified RpHLuorin2 (Figure 1; Supplementary Figure 1), we observed for GPI-pHluorin2 that the ratio of fluorescence with 405 and 470 nm excitations changed over a narrower range of pH values (5.5–7.5; Supplementary Figure 3a) than with FLIM imaging (5.5–7.5; Figure 2). For the MGAT2 and GalT probes, we observed similar pH values as with the FLIM-based measurements (Supplementary Figure 3b; MGAT2 pH 6.5 (95%CI ± 0.16), GalT pH 6.1 (95%CI ± 0.13)). However, the spread of the data was considerably (~2-fold) larger for the ratiometric approach. As a result, less cells have to be analyzed with FLIM-based approach to accurately determine the pH. To illustrate this point, we performed Bootstrap statistical analysis where we sampled our datasets to estimate the 95% CI based on samples of increasing numbers of cells (Supplementary Figure 4). Based on this analysis, we estimate that for the FLIM-based approach >16 cells needed to be measured for an accurate estimation of the pH for the ER, MGAT2, GalT and LAMP1 markers. However, for the ratiometric approach, approximately 2-fold more cells needed to be analyzed to reach a similar 95%CI.

To further characterize the RpHLuorin2 FLIM system, we challenged cells with the vacuolar H^+^-ATPase (V-ATPase) inhibitor Bafilomycin A1 (BafA1)^53^. The mammalian V-ATPase is a protein pump that acidifies intraorganellar lumina by translocating protons across the membrane^54,55^. Our experiments with the MGAT2 and GalT probes showed that challenging the cells for 1 hr with 200 nM BafA1 resulted in a reduced acidification of both the cis- and *trans*-Golgi apparatus, although this perturbation was incomplete as the pH did not reach completely neutral values (Figure 4). Taken together, our data show that the RpHLuorin2 FLIM system is highly suitable for intracellular pH measurements with only a single image acquisition.

**Figure 4.**
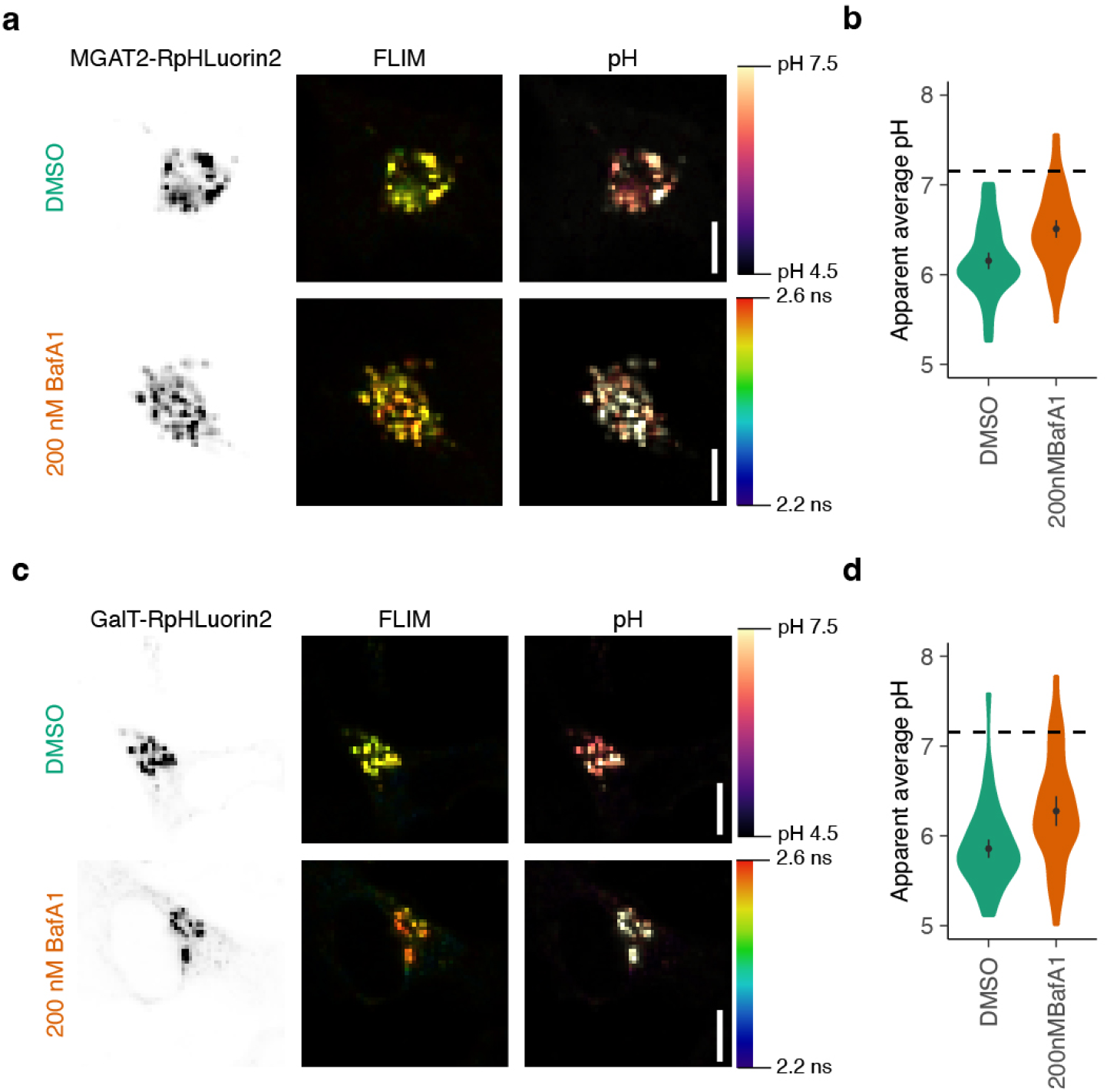
Incomplete blockage of Golgi acidification by Bafilomycin A1. (a) Representative confocal micrographs of HeLa cells expressing MGAT2-RpHLuorin2 incubated for 1 hr in the absence (solvent control DMSO) or presence of Bafilomycin A1 (200 nM BafA1). To generate the FLIM (fluorescence lifetime imaging microscopy) images (middle column), the intensity images (left column) were convoluted with the fluorescent lifetime value per pixel and pseudo-colored. To generate the pH images, the lifetimes were converted to the calculated pH per pixel and also convoluted with the fluorescence intensities (right column). Scalebars, 10 μm. (b) Quantification of average pH values from panel (a). N = 72 (DMSO) and 77 (BafA1) cells from 4 independent experiments. The dashed lines indicate the average pH of the ER from Figure 3c. (c-d) Same as panels (a-b), but now for GalT-RpHLuorin2. N = 68 (DMSO) and 50 (BafA1A) cells from 4 independent experiments.

### Dynamic measurements of pH in the secretory pathway

To evaluate whether our method would be able to measure dynamic changes in pH, we started by measuring the pH of the medial-Golgi marker MGAT2-pHLuorin2 in the presence of fungal metabolite Brefeldin A (BFA). BFA is a potent inhibitor of ER-Golgi trafficking and causes the relocation of Golgi-resident enzymes to the ER^56,57^. We therefore expected a substantial increase in pH when MGAT2-RpHLuorin2 expressing cells were challenged with BFA. Indeed, we measured an apparent average pH of 7.1 (95%CI ± 0.07) in the BFA-challenged cells compared to an apparent average pH of 6.4 (95%CI ± 0.08) in the vehicle control cells (Figure 5a, b). This result means that our system is capable of measuring dynamic alterations of pH in living cells.

**Figure 5.**
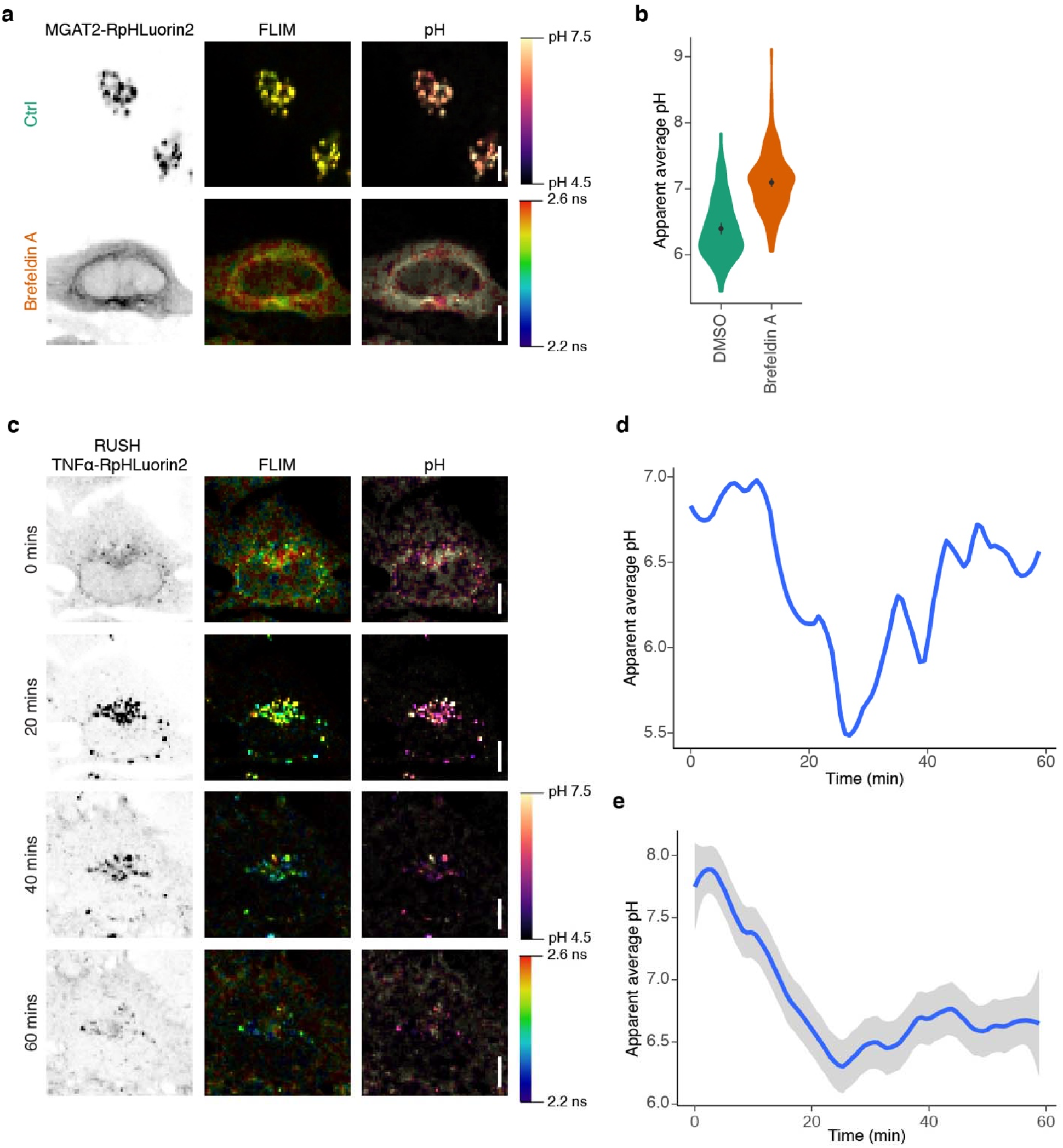
Dynamic pH measurements along the secretory pathway. (a) Representative confocal micrographs of HeLa cells expressing MGAT2-RpHLuorin2 in the absence (Ctrl, green) or presence of Brefeldin A (orange). The intensity image (left column) was convoluted with the fluorescent lifetime value per pixel and pseudo-colored (middle column). The intensity image was also convoluted with the calculated pH per pixel and pseudo-colored (right column). FLIM, fluorescence lifetime imaging microscopy. Scalebars, 10 μm. (b) Quantification of average pH values from panel (a). N = 110 (DMSO) and 165 (Brefeldin A) cells from 2 – 3 independent experiments. (c) Representative confocal micrographs of HeLa cells expressing RUSH TNFα-RpHLuorin2 in the absence of biotin (0 min) or 20, 40 and 60 minutes after biotin addition. The intensity image (left column) was convoluted with the fluorescent lifetime value per pixel and pseudo-colored (middle column). The intensity image was also convoluted with the calculated pH per pixel and pseudo-colored (right column). FLIM, fluorescence lifetime imaging microscopy. Scalebars, 10 μm. See also Supplementary Movie 1. (d) Quantification of average pH values of the cell shown in panel (c) and Supplementary Movie 1. (e) Average pH measured of all cells expressing RUSH TNFα-RpHLuorin2. N = 29 from 2 independent experiments.

Next, we employed FLIM-based measurements to monitor the changes of the pH in real-time along the secretory pathway. To this end, we chose the secreted cytokine tumor necrosis factor alpha (TNF-α) as a model protein that is transported through the secretory pathway. Using the Retention Using Selective Hooks (RUSH) system^51^, we synchronized the transit of TNF-α along the secretory pathway. RUSH uses the expression of two separate constructs in the cell: (i) the hook construct, which is an ER-targeting sequence fused to streptavidin, and (ii) the reporter construct which is the protein of interest (i.e., TNF-α) fused in tandem to a streptavidin binding protein (SBP) and a fluorescent protein (RpHLuorin2). When biotin is absent from the culture medium, the reporter construct is held at the ER through an interaction of streptavidin of the hook construct and the SBP. When biotin is added to the culture medium, biotin outcompetes this interaction and the reporter construct is released and transits along the secretory pathway in a synchronized fashion.

In our case, we used the KDEL-motif as a targeting sequence for the ER^51^, and used a TNFα-SBP-RpHLuorin2 fusion protein (RUSH TNFα-SBP-RpHLuorin2) as the reporter construct, so that we could follow the dynamic transit of TNF-α from the ER to the plasma membrane (Figure 5c-e, Supplementary Movie 1). We confirmed the subcellular localizations with immunolabeling experiments where we fixed cells expressing TNFα-SBP-RpHLuorin2 at discrete time intervals after biotin addition and immunolabeled for organellar markers for the ER (PDI), Golgi network (GM130) and plasma membrane (WGA) (Supplementary Figure 5).

In the absence of biotin in the cell culture medium, when all the TNFα-SBP-RpHLuorin2 reporter construct was trapped within the ER, we measured an apparent average pH of 7.58 (95%CI ± 0.46) (Figure 5c-e, Supplementary Movie 1). In the ~25 min following the addition of biotin to the cells, TNFα-SBP-RpHLuorin2 was trafficked through the Golgi and the apparent average pH gradually decreased to around pH 6. At later time points, the pH gradually increased again as more TNFα-SBP-RpHLuorin2 reached the plasma membrane. As HeLa cells express the protease TACE, TNF-α likely dissociates from the plasma membrane^58–60^.

After biotin addition, the TNF-α-RpHLuorin2 became concentrated in the Golgi network, leading to a local increase of signal at this position. In order to not saturate the signal at this timepoint, we had to use a low intensity of excitation light for the RUSH experiments. However, at the start of the experiments (i.e., prior to biotin addition), the TNF-α-RpHLuorin2 construct was located at the ER, which in mammalian cells is diffuse and scattered through the entire cytoplasm. Likely as a consequence, the photon count/pixel at earlier timepoints was low leading to an overestimation of the pH particularly for the ER localization. Also because of the limited number of photons, we fitted the fluorescence lifetime histograms with a single exponential decay function and report the apparent average pH per cell^61^.

This result demonstrates that FLIM-based pH measurements are a suitable method to determine intraorganellar pH with high temporal resolution.

## Discussion

In this study, we measured the pH in various subcellular compartments using FLIM of the pH-sensitive fluorescent protein RpHLuorin2. Consistent with previous literature, we observed a clear acidification of luminal pH through the secretory pathway^1–3^. The fusion of RpHLuorin2 is not restricted to the proteins we described here; this system is applicable to any other intraorganellar measurement, provided RpHLuorin2 can be fused to a luminal domain of a protein residing in the target organelle. Furthermore, additional applications include combining RpHLuorin2 with other fluorescence (lifetime) based probes to measure pH and other cellular processes simultaneously within the same cell.

We also show that the FLIM approach enables to measure pH with a kinetic resolution high enough to follow the dynamic transit of a cargo molecule along the secretory pathway. In this respect, our approach complements measurements of the exocytic pathway using Vero and Shiga toxins labeled with-pH sensitive fluorophores^31^. These toxins are endocytosed by receptor-mediated endocytosis and then transit via the Golgi network to the ER, a process which can be followed by microscopy. While this approach also enables to measure pH along the secretory pathway, a disadvantage is that it does not allow to follow the pH of designated secretory targets. For example, after transit through the Golgi, TNF-α is reported to traffic via designated sub-compartments of recycling endosomes to the plasma membrane^62^, and it is not known whether Shiga toxins also traffic via these compartments.

Compared to excitation-based ratiometric imaging, the key improvement of our study is the usage of FLIM. Ratiometric imaging of pHLuorin and derivatives^30,34,35^ requires the sequential recording of the fluorescent protein at both 405 and 470 nm excitation wavelengths, while the emission is recorded at the same wavelength. Although certain optical schemes such as split-beam excitation might facilitate fast switching between excitation wavelengths, this sequential excitation intrinsically limits the temporal resolution and consequently limits the applicability for pH determination in dynamically moving and reshaping organelles. FLIM mitigates this issue, as only a single recording with a single excitation wavelength is required. Moreover, FLIM measurements are not dependent on laser intensity^41,42^, while ratiometric measurements can easily be affected by fluctuations in excitation laser power. FLIM measurements are thus more comparable between experiments, as supported by our findings that the spread in the data is larger for the ratiometric than for the FLIM approach. Another advantage of the FLIM-based approach is that reference measurements can be used over independent experiments, because the fluorescence lifetime is independent of the fluorescence intensity^41,42^. This is an advantage over the ratiometric approach, where small differences in the laser intensity and alignment of the microscope can have a major effect. However, a disadvantage is that it requires access to a FLIM microscope, whereas ratiometric imaging can be performed on most confocal and epifluorescence microscopes.

In contrast to another study which relies on equilibrating pH with the ionophore monensin^30^, we used GPI-anchored RpHLuorin2 to obtain calibration curves with defined pH buffers, because monensin is a known inhibitor of physiological Golgi transport, thereby likely affecting the observed fluorescence lifetime values^63–69^.

Defects in the regulation of pH are a hallmark of a wide range of disease, including disorders of glycosylation^4,19,21–25^, cancer^70^, neurodegenerative diseases^71–74^, mitochondrial disorders^75^ and lysosomal storage disorders^76^. The tools we presented in this study offer a method to assess intraorganellar pH using FLIM. Our data show that FLIM is more accurate than ratiometric imaging. Moreover, due to its high temporal resolution, it not only enables to measure pH in static compartments, but also to measure the dynamic changes that a protein experiences during its trafficking along the secretory pathway.

## Acknowledgments

We thank the following people for constructs: Hesso Farhan and Franck Perez (Str-KDEL_ManII-SBP-EGFP; Addgene plasmid #65252, Str-KDEL_TNF-SBP-EGFP; Addgene plasmid #65278), Lei Lu (piRFP670-N1-GalT; Addgene plasmid #87325), Carsten Schultz and André Nadler (GPI-mRFP), and Esteban Dell’Angelica (LAMP1-mGFP, Addgene plasmid #34831). We thank the Microscopy Imaging Center of the Radboud Institute for Molecular Life Sciences for use of their microscopy facilities. G.v.d.B. is funded by a Young Investigator Grant from the Human Frontier Science Program (HFSP; RGY0080/2018) and a Vidi grant from the Netherlands Organisation for Scientific Research (NWO-ALW VIDI 864.14.001). G.v.d.B has also received funding from the European Research Council (ERC) under the European Union’s Horizon 2020 research and innovation program (grant agreement No. 862137).

## Author Contributions

P.T.A.L., M.t.B., and G.v.d.B. designed and performed the experiments and wrote the paper. M.I. performed the ratiometric pH measurements in HeLa cells and contributed to the writing of the paper.

## Methods

### Cloning

The sequence of RpHLuorin2^35^ was synthesized by Genscript for both recombinant (codon optimized for *E. coli*) and mammalian cell expression. Synthetic RpHLuorin2 codon optimized for *E. coli* for recombinant protein production was inserted in pET-28a(+) (EMD Biosciences) with restriction sites NdeI and XhoI. The construct for cytosolic expression of RpHLuorin2 was generated by replacing EGFP in pEGFP-N1 (Clontech) with synthesized RpHLuorin2 using restriction sites AgeI and BsrGI. ER-targeting of RpHLuorin2 was achieved by inserting the signal sequence of calreticulin (MLLSVPLLLGLLGLAVA) and flexible GGSGGS linker before RpHLuorin2 and adding a KDEL motif for luminal ER retention after RpHLuorin2. Targeting to the MGAT2-positive compartment was achieved by inserting synthetic truncated MGAT2 (residues 1-89 of Uniprot Q10469, Genscript) in the vector for cytosolic expression of RpHLuorin2 with restriction sites EcoRI and BamHI. The starting codon of RpHLuorin2 was removed and a flexible GGSGGS linker was added between the two protein fragments. Targeting to the GalT-positive compartment was achieved by inserting truncated GalT (Addgene plasmid #87325) in the vector for cytosolic expression of RpHLuorin2 with restriction sites SmaI and BamHI. The starting codon of RpHLuorin2 was removed and a flexible GGSGGS linker was added between the two protein fragments. Targeting to LAMP1-positive compartments was achieved by inserting the signal sequence of LAMP1 (LAMP1-mGFP^77^, Addgene plasmid #34831) N-terminal to RpHLuorin2 with restriction sites HindIII and AgeI. Then, the luminal domain of LAMP1 was placed C-terminal to RpHLuorin2 after a flexible GSGS linker with restriction sites BsrGI and NotI. GPI-RpHLuorin2 was generated by replacing mRFP from GPI-mRFP with synthetic RpHLuorin2 with restriction sites XmaI and NotI. RUSH TNFα-RpHLuorin2 was generated by replacing EGFP from Str-KDEL_TNF-SBP-EGFP (Addgene plasmid #65278) with synthetic RpHLuorin2 (Genscript) with restriction sites SbfI and BsrGI. All sequences were verified by Sanger sequencing prior to transfection. All generated plasmids from this study have been deposited at Addgene.

### Cell culture and transfection

HeLa cells (authenticated by ATCC through their human STR profiling cell authentication service) were maintained in high glucose DMEM with Glutamax (Gibco 31966021), supplemented with 10% fetal calf serum (FCS, Greiner Bio-one, Kremsmünster, Austria) and antibiotic-antimycotic solution (Gibco 15240-062). Cells were regularly tested for mycoplasma contamination. HeLa cells were transfected with plasmid vectors using Fugene HD (Promega E2311), using the recommended protocol of the manufacturer. Cells were imaged 48 hours post-transfection. Only cells expressing low to moderate levels of the transfected plasmids, based on fluorescence intensity and manual localization scoring, were chosen for subsequent microscopic analyses.

### pH calibration buffers

Buffers with defined pH for the generation of a calibration curve were prepared as described previously^30^. Calibration buffers contain 125 mM KCl, 25 mM NaCl, and 25 mM N-[2-hydroxyethyl]-piperazine-N-[2-ethanesulfonic acid] (HEPES, pH 7.5 or 7.0) or 25 mM 2-[N-morpholino] ethanesulfonic acid (MES, pH 6.5, 6.0, 5.5, 5.0 or 4.5). Each buffer solution was adjusted to the appropriate final pH using 1 M NaOH or 1 M HCl at 37°C.

### Recombinant protein expression and purification

RpHLuorin2 in pET-28(a)+ vector was transformed in BL21(DE3) *E. coli*. Cells were grown in 2x yeast extract - tryptone medium and induced with 250 μg/mL β-D-1-thiogalactopyranoside (IPTG) at an OD600 of 0.8 for 2 hrs at 37°C and 200 rpm. Cells were pelleted at 3000 × *g* at 4°C for 30 mins and subsequently lysed with B-PER (Thermo Scientific) supplemented with 50 U DNAseI, 1:500 lysozyme and protease inhibitor cocktail (Roche). Lysates were then cleared by centrifugation (20,000 × *g* at 4°C for 10 mins) and supernatants with recombinant protein were loaded onto nickel – nitrilotriacetic acid (Ni-NTA) bead columns. Ni-NTA beads were then washed five times with 10 mM Tris-HCl pH 7.6 and 150 mM KCl, and proteins were eluted in 10 mM Tris-HCl pH 7.6, 150 mM KCl and 200 mM imidazole. To remove the elution buffer, purified protein was dialyzed overnight against ddH_2_O using 2 mL Slide-A-Lyzer 10K MWCO tubes (Thermo Scientific). Protein concentration was determined using the Micro BCA Protein Assay Kit (Thermo Scientific). Fluorescence of recombinant pHLuorin2 in each pH calibration buffer was measured using Eppendorf semimicro Vis Cuvettes on an LS 55 Fluorescence spectrometer (PerkinElmer). A xenon lamp was used as the excitation source. Samples were excited at 405 and 470 nm, and the emission was recorded at 508 nm.

### Confocal microscopy

Imaging of cells took place in Leibovitz’s L-15 medium, with the exception of the samples for the pH calibration curve and the recombinant protein measurements. GPI-RpHLuorin2 and recombinant RpHLuorin2 (10 μM) measurements were performed in the pH buffers mentioned above. For calibration with GPI-RpHLuorin2, cells were preincubated in pH buffer for 15 mins at 37°C to achieve sufficient pH equilibration. For the Bafilomycin A1 assay, cells were treated with 200 nM Bafilomycin A1 (Cayman Chemicals) in DMSO for 1 hour prior to live imaging. For the Brefeldin A assay, cells were treated with 10 μg/mL Brefeldin A (Cayman Chemicals) in DMSO for 15 mins prior to live imaging. Imaging was performed on a Leica SP8 SMD system at 37°C, equipped with an HC PL APO CS2 63x/1.20 WATER objective. pHLuorin2 was excited at 488 nm with a pulsed white light laser, operating at 80 MHz. Photons were collected for one minute or 30 seconds for time-lapse experiments with a HyD detector set at 502 – 530 nm and lifetime histograms of the donor fluorophore were fitted with a monoexponential decay function convoluted with the microscope instrument response function in Leica LAS X. For reconstructing the images, tiff files with τ values were generated using FLIMFit^78^ and 2 x 2 spatial binning, and then convoluted with the fluorescent intensities using a custom-written ImageJ macro. Ratiometric pH measurements were done similarly to the FLIM measurements, but instead the imaging was performed on a Leica SP8 SMD system at 37°C, equipped with an HC PL APO CS2 63x/1.20 Water objective or a Zeiss LSM800 system at 37°C, equipped with a Plan Apochromat 1.4x Oil objective. RpHLuorin2 was excited at 405 nm and 488 nm sequentially, images were acquired with an emission wavelength bandwidth (495–560 nm) that included the emission wavelength of 508 nm.

### Colocalization analysis

25.000 were plated on cleaned 12 mm glass coverslips (Electron Microscopy Services), transfected as described above with RUSH TNFα-RpHLuorin2. The RUSH assay was performed after 48 hours, and cells were fixed with 4% paraformaldehyde for 15 mins at room temperature at the indicated times and quenched with 50 mM NH_4_Cl in PBS for 10 mins. Wheat germ agglutinin (WGA) staining was performed without permeabilization and before any other staining step. Briefly, cells were washed with Carbo-Free Blocking Solution (Vector Laboratories, supplemented with 1 mM MgCl_2_ and CaCl_2_) and incubated with 4 μg/mL biotinylated WGA (Vector Laboratories) diluted in Carbo-Free Blocking Solution. Cells were then incubated with Strepatividin-Alexa Fluor 647 (ThermoScientific). Anti-GM130, anti-PDI and anti-GFP (recognizes RpHLuorin2) was performed with permeabilization. Cells were permeabilized and blocked in 2% normal donkey serum (Rockland) and 0.1% saponin (permeabilization buffer) for 30 mins at RT. Primary antibodies were diluted in permeabilization buffer and incubated for 1 h at room temperature: mouse anti-PDI (clone RL90, Novus Biologicals), 10 μg/mL; mouse anti-GM130 (clone 35/GM130, BD Biosciences), 2.5 μg/mL, and rabbit anti-GFP (Rockland), 20 μg/mL. Secondary antibodies were also diluted in permeabilization buffer and incubated for 1 h at room temperature: donkey-anti-rabbit Alexa Fluor 488 and donkey-anti-mouse Alexa Fluor 647 (both ThermoFisher, 5 μg/mL). Cells were finally washed with 0.1% Triton X-100 in PBS to remove any background staining and mounted usin mounting medium with 0.01% Trolox (6-hydroxy-2,5,7,8-tetramethylchroman-2-carboxylic acid) and 68% glycerol in 200⍰mM sodium phosphate buffer at pH 7.5 with 0.1⍰μg/mL DAPI (4⍰,6-diamidino-2-phenylindole). Coverslips were sealed with nail polish. Cells were imaged on a Leica SP8 SMD confocal laser scanning microscope, equipped with an HC PL APO CS2 63x/1.20 WATER objective. Colocalization analysis was performed using the *pearsonr* function from the Python package SciPy^79^. Individual cells were first saved to separate .tiff files with ImageJ, and then processed in a fully automated and unbiased fashion using the *pearsonr* function.

### Quantification and statistical analysis

All mean values represent the average of all cells analyzed. pH calibration curves were fitted with an emperical dose response function as described by equation (1) with the *MM.3* function from the R package *drc*^80^.

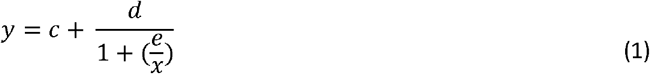

Where *y* is the fluorescence intensity or lifetime, *x* is the pH and *c, d* and *e* are fit parameters.

Bootstrap analyses were performed using R statistical software using the interpolated pH values from all cells measured in the lifetime and ratiometric measurements. Briefly, 95% confidence intervals were determined for random samples of measurements (starting from n=2, up to the number of cells per condition). This process was repeated for 10,000 iterations using the *replicate* function.

All comparisons were first checked for similar mean and median values and acceptable (< 3x) difference in variance. Statistical analysis of three or more groups was performed using a one-way ANOVA, followed by a post-hoc Tukey’s honestly significant difference test. p < 0.05 was considered significant. *p < 0.05, **p < 0.01, ***p < 0.001, ****p ≤ 0.0001. All statistical analyses were performed using R statistical software. All numerical data were visualized using R package *ggplot2*^81^, with violins representing the overall distribution of the data and means ± 95% CI overlaid.

### Data and code availability

All raw data, including R scripts and ImageJ macros, have been deposited to Zenodo.

