## Supplementary Information for "Fluorescence lifetime imaging of pH along the secretory pathway"

**Supporting Information**

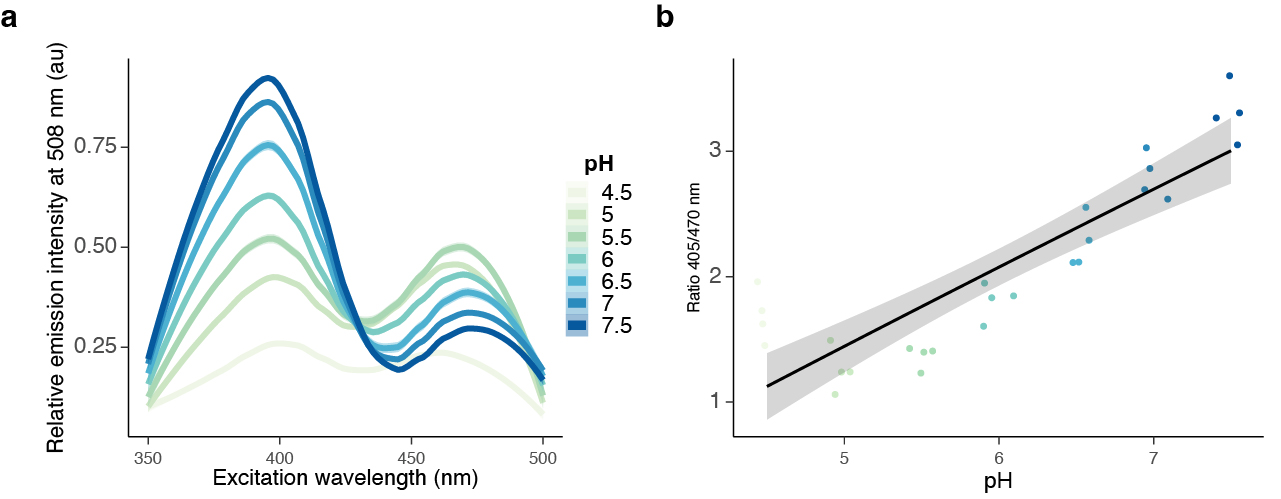

**Supplementary Figure 1. Fluorescence excitation spectra of recombinant RpHLuorin2.**

(a) Representative fluorescence excitation spectra (emission 508 nm) of recombinant RpHLuorin2 in buffers of defined pH were collected with a fluorescent spectrometer.
(b) Calibration curve from panel (a). The ratios of fluorescence at 405 over 470 nm were plotted as a function of pH and fitted with a three-parameter Michaelis-Menten fit. Data from four independent experiments.

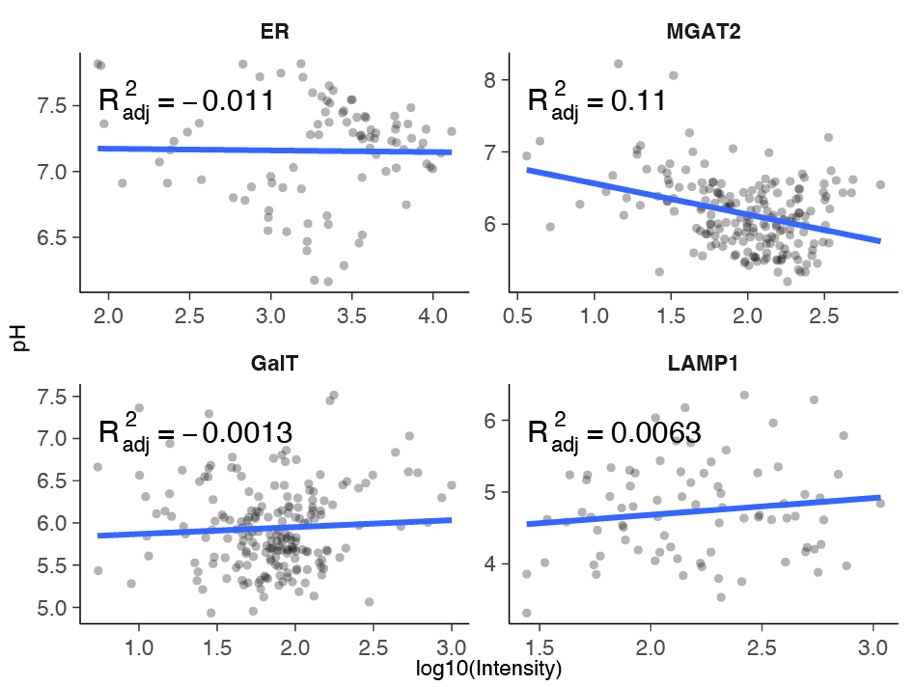

**Supplementary Figure 2: FLIM-based pH measurements are independent of expression intensity.**

Correlation analysis of the images analyzed in main Figure 3. Per marker the pH is plotted on the y-axis and the log10 of the fluorescent intensity on the x-axis. No linear correlation was observed between pH and fluorescent intensity.

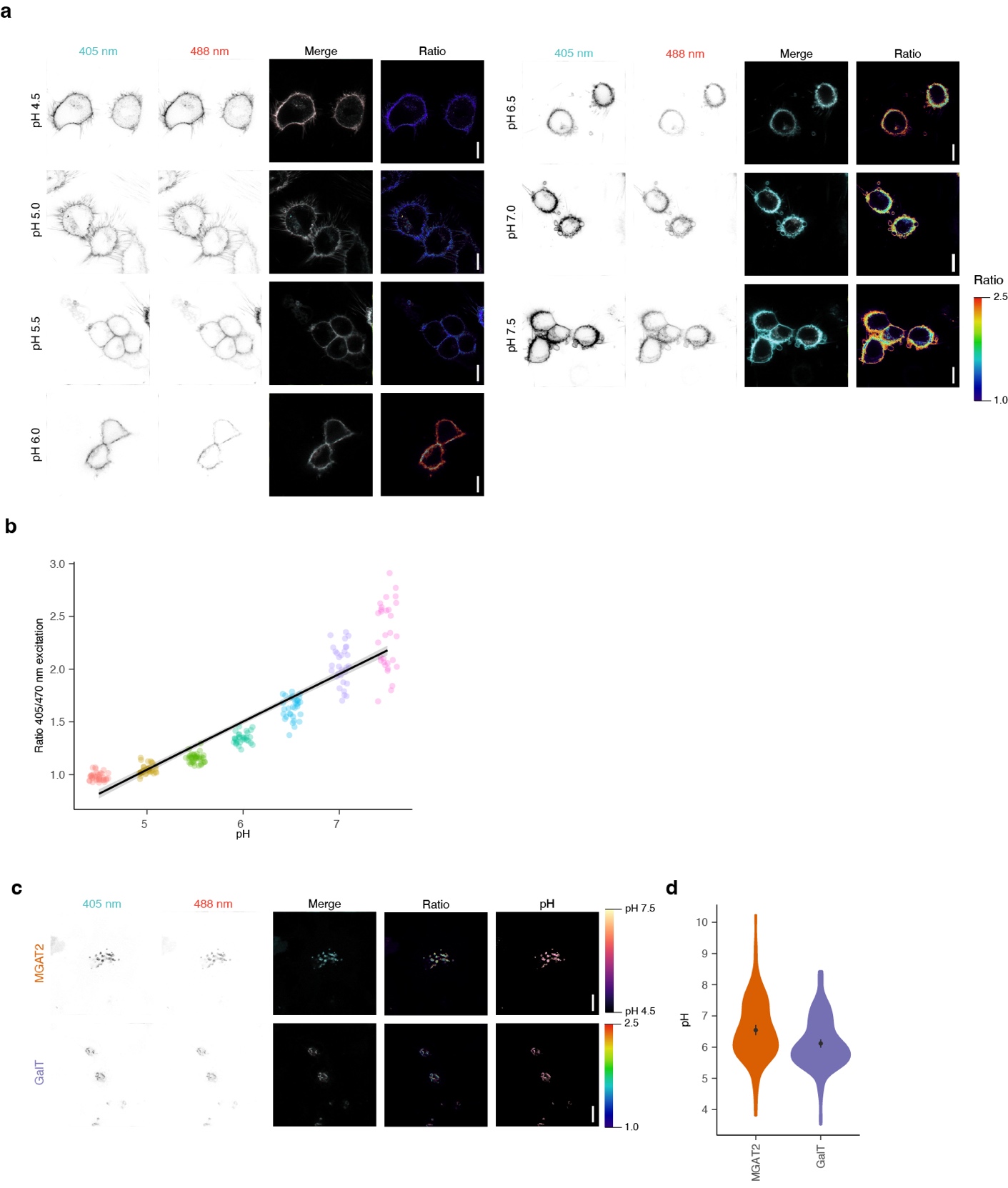

**Supplementary Figure 3. Measurements of organellar pH using ratiometric imaging of RpHLuorin 2.**

(a) Representative confocal micrographs of HeLa cells expressing GPI-RpHLuorin2 in defined calibration buffers. 405 nm excitation in magenta, 470 nm excitation in green. The ratio image (right column) was calculated by dividing the 405 nm excitation channel by the 470 nm excitation channel and pseudo-colored. Scalebars, 10 µm.
(b) pH dependence of HeLa cells expressing GPI-RpHLuorin2 in defined pH calibration buffers from the ratiometric images of panel (a). Data from one representative experiment is shown as the ratio values between experiments are not directly comparable. For this experiment: N = 37 (pH 4.5), 34 (pH 5.0), 39 (pH 5.5), 30 (pH 6.0), 32 (pH 6.5), 30 (pH 7.0), and 30 (pH 7.5).
(c) Representative confocal micrographs of HeLa cells expressing the mentioned RpHLuorin2 fusion constructs. The ratio image (right column) was calculated by dividing the 405 nm excitation channel by the 470 nm excitation channel and pseudo-colored. Scalebars, 10 µm.
(d) Quantification of average pH from panel (c). N = 138 (MGAT2) and 151 (GalT) from 3 independent experiments.

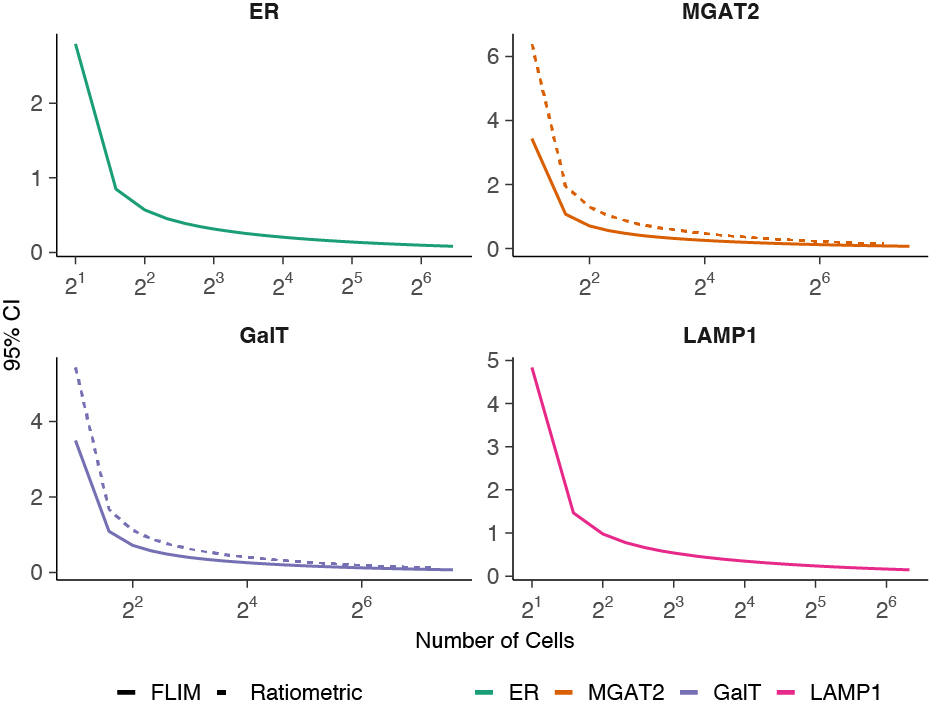

**Supplementary Figure 4. FLIM-based pH measurements are more accurate than ratiometric pH measurements.**

Bootstrap analysis of sample variance between FLIM and ratiometric pH measurements. Incremental numbers of cells were randomly sampled from the data of Figure 3 (FLIM; solid curves) and Supplementary Figure 3 (ratiometric; dashed curves for MGAT2 and GalT). The 95% CI (confidence interval) was calculated and the mean for 10,000 repetitions was plotted.

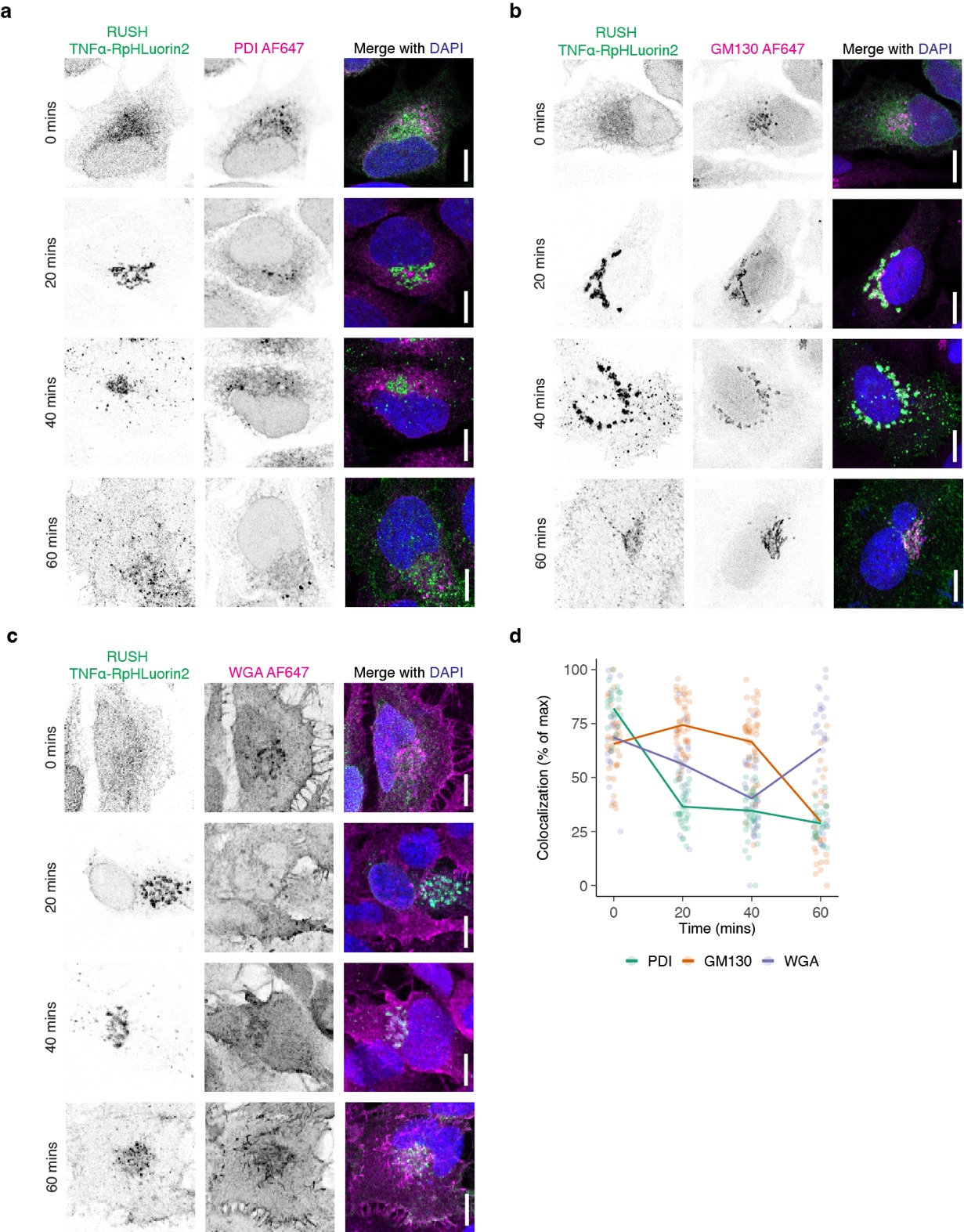

**Supplementary Figure 5. Colocalization of TNFa-SBP-RpHLuorin2 with organellar marker proteins.** Representative confocal micrographs of HeLa cells expressing RUSH TNFa-SBP-RpHLuorin2 (green in merge). The cells were cultured with biotin for the mentioned times, fixed in paraformaldehyde, and immunolabeled for the ER marker PDI (panel (a); magenta in merge), the Golgi marker protein GM130 (panel (b)), or the plasma membrane marker WGA (wheat germ agglutinin, without permeabilization, panel (c)). Blue: DAPI. Scalebars, 10 µm**.**

(d) Quantification of colocalization from panels a-c. Pearson correlation coefficients were normalized between 0 and 1 per marker for all cells, then multiplied by 100 to get the percentage.

**Supplementary Movie 1.** Time-lapse FLIM imaging of RUSH TNFα-RpHLuorin2 in HeLa cells after biotin addition. The left panel shows the fluorescent intensity, the middle panel the fluorescent intensity convoluted with the lifetime, and the right panel the fluorescent intensity convoluted with pH values.
